# RNA sequencing identifies dysregulated circular RNAs in early-stage breast cancer

**DOI:** 10.1101/506246

**Authors:** Vittal Rangan Arvinden, Arunagiri Kuha Deva Magendhra Rao, Balaiah Meenakumari, Priya Ramanathan, Shirley Sundersingh, Velusami Sridevi, Thangarajan Rajkumar, Zdenko Herceg, Samson Mani

## Abstract

**Background:** Breast cancer is a major cause of cancer related death in women worldwide. Molecular diagnostic markers that are detectable in early-stage breast cancer can aid in effective clinical intervention. Circular RNAs are a recently identified group of non-coding RNA with potential role in cancer development and progression. In this study, we aimed to identify circular RNAs specific for early stage breast cancer.

**Method:** Circular RNA expression profile was analyzed in early-stage breast cancer tissues (N=5), matched normal counterparts (N=5) and absolute normal samples (N=5) by RNA-sequencing that enables a comprehensive analysis of RNA expression across the transcriptome. Two different algorithms, find_circ and DCC were used to identify the differentially expressed circular RNAs.

**Results:** A total of 58 and 87 circular RNAs were found to be differentially expressed by find_circ and DCC algorithms, respectively, among which 26 circular RNAs were common. Hsa_circ_0001946 (CDR1-as) was found to be upregulated in early stage breast cancer along with other novel circular RNAs (hsa_circ_0008225, hsa_circ_0007766, hsa_circ_0016601). We also found that a few of the identified circular RNAs harbor microRNA binding sites which can lead to microRNA sponging activity and pre-microRNA sequences which can generate mature microRNAs. The identified circular RNAs that are differentially regulated in early stage breast cancer can be of potential diagnostic/prognostic importance.

**Conclusion:** Circular RNA are differentially expressed in the early-stage breast cancer with potential application in early diagnosis and prognosis. The differentially expressed circular RNA can sequester microRNA and can act as microRNA precursor as well.

## INTRODUCTION

Breast cancer is the leading cause of cancer related deaths among women with 1.6 million new cases and 0.52 million deaths each year **^1^**. Early detection can aid in better treatment outcomes. In recent years, non-coding RNA comprising different types of small and long non-coding RNA have shown to be dysregulated in cancer with accumulating evidence in diagnosis and prognosis **^2-5^**. Circular RNAs (circRNA) are a large group of emerging non-coding RNA that result from back splicing of linear RNA. Circular RNAs are covalently closed circular single stranded RNA molecules lacking both 5’-cap and 3’-tail **^6^**. Classified under the group of non-coding RNA, it is interesting to note that circular RNAs with ORF and IRES sequence can also code for peptides and therefore are a hybrid set of RNA with both structural and functional importance **^7^**. High-throughput sequencing and algorithms like find_circ, CIRCexplorer, CIRI, DCC etc have aided in emerging number of circular RNAs. Few databases like circRNADb and circBase have listed 32914 (exonic) and 92375 circular RNAs respectively reported from various studies **^8,9^**. Circular RNA contain a combination of exon or intron or an untranslated region of a linear RNA. The most common spliced-out forms of circular RNAs carry exonic sequences of the parental transcript. But those circular RNAs harboring UTR sequence and with intronic features may possess regulatory function. The differentially expressed circular RNAs are attributable to molecular events in carcinogenesis. Circular RNAs are reported to be deregulated in hepatocellular carcinoma, esophageal squamous cell carcinoma, colorectal cancer, bladder cancer and breast cancer**^10,11^**. A recent study found, circGFRA1_5-7 overexpression in estrogen receptor positive (ER+) breast tumors while circ_RPPH1 and circ_ERBB2_7-11 were abundantly expressed in human epidermal growth factor receptor 2 positive (HER2+) breast tumors samples but their role corresponding to the subtypes are yet to be elucidated **^12^**. Similarly circ-FOXO3 was observed to be downregulated in human breast cancer tissues while its ectopic expression induced apoptosis in mouse xenografts^13^. Circ_103110, circ_104689 and circ_104821 were previously reported to be more of diagnostic use in infiltrating ductal carcinoma **^14^**. Moreover, circular RNAs are abundantly found in bodily fluids like saliva, serum and also sorted into exosomes **^11,15,16^**. Hence, detectable circular RNAs specific to early stage breast cancer can be of immense importance in diagnosis and prognosis.

Understanding about the functional role of circular RNAs in tumorigenesis and progression is still at its early stage. The structure of circular RNA provides a longer half-life, increasing the probability of interaction with other biomolecules **^17^**. Most common role as competing endogenous RNAs is a well-known function of circular RNA. Circular RNA *CDR1-as* through its sponging activity was shown to hamper miR-7 mediated gene regulation *CDR1-as* apart from sponging, can act as buffer by sustained release of miR-7 or as a reservoir when miR-671 cleaves *CDR1-as* using RISC assembly to mediate control of targets **^18^**. The miR7-*CDR1-as* regulation has been associated with hepatocellular carcinoma and other disorders **^19^**. Various circular RNAs have been shown to repress microRNA, altering its function in breast cancer. Hsa_circ_0001982, circABCB10 and circRAK3 were reported to be sponging miR-143, miR-1271 and miR-3607 respectively leading to breast carcinogenesis and metastasis **^20-22^**. Nevertheless, newly identified breast cancer specific circular RNA will disclose potential circular RNA-microRNA relationship uncovering unknown underlying ncRNA interactions contributing to breast carcinogenesis.

In this study, we have attempted to identify circular RNAs differentially expressed in early stage breast cancer using RNA-sequencing. The assessment of potential circular RNA-microRNA interaction revealed multiple binding sites for microRNA, suggesting that the sponging of microRNA or Argonaute mediated degradation of circular RNA may be the dual outcome of these interactions. In addition, we have also explored the function of circular RNAs harboring sequences with the potential to yield mature microRNA. Overall we have identified aberrantly expressed circular RNAs in early stage breast cancer, which may be used as stable diagnostic and prognostic biomarkers based on further evaluation.

## MATERIALS AND METHODS

### Study samples

Tissues were surgical specimen obtained from tumor bank at Cancer Institute (WIA), Chennai, India. Breast cancer cases of stage I-IIA were selected for the study. Tumor tissue (N=5) sections were histo-pathologically confirmed to contain more than 70% of tumor cells. Adjacent normal breast tissue from the same patient were used as matched normal samples (N=5). From patients undergoing surgery for non-malignant breast conditions, tissues samples distally away from the pathological indication within the breast were used as normal samples (N=3) (Supplementary Table 3). All the surgical specimens were obtained after informed consent from the participants. The study was conducted in accordance with the Declaration of Helsinki, and the protocol was approved by the Cancer Institute Ethical Committee, Cancer Institute (WIA), on 24-10-2013 (Project title: Identification and functional characterization of non-coding RNAs (lncRNA/microRNA) in early stage breast cancer). The RNA sequencing data was deposited in SRA (Accession ID: SRP156355).

### RNA-sequencing

Total RNA isolated from frozen tissues was depleted for ribosomal RNA using the Ribo-Zero Gold rRNA removal kit (Illumina, USA). The cDNA libraries were prepared using TruSeq Stranded Total RNA Library Prep kit (Illumina, USA) following standard protocol. Quality profile of library was assessed using Bioanalyzer 2100 (Agilent, Santa Clara, CA) and 100 bp PE sequencing was carried out by HiSeq 2000 (Illumina, San Diego, CA).

### Identification of circular RNAs

FASTQ files were mapped against human reference genome (hg38) using bowtie2. The unmapped reads from BAM files were used as input for find_circ algorithm ^46^. Briefly, the 20-mer sequence from both ends of the unmapped reads were aligned against the anchor position within the spliced exon. BED files were filtered for circular RNAs with unambiguous breakpoints carrying characteristic GT/AG splice sites and a head-to-tail reverse alignment of the anchors. These circular RNAs obtained were filtered by 5 unique back spliced reads in at least one sample and ≥ 2 reads in minimum of 2 out of other biological replicates of each group ^47^.

We used another algorithm DCC developed to detect circular RNAs using output from STAR read mapper and refine data with series of inbuilt filters ^48,49^. Reads were mapped to hg38 reference genome and ENSEMBL GrCh.38.37 GTF file using STAR alignment tool to obtain information on chimeric splice junctions. The generated ‘chimeric.out.junction’ file contains chimerically aligned reads including circular RNA junction spanning reads. Circular RNAs identified by both tools were then annotated using circus R package using UCSC hg38 TxDB. The differential expression of circular RNAs detected by both find_circ and DCC was carried out using DESeq2 R package. Circular RNAs with |fold change| ≥ 2 and p-value ≤ 0.05 were considered to be differentially expressed between sample group.

### MicroRNA – circular RNA interaction prediction

MicroRNA target binding sites of circular RNAs were predicted using *CircInteractome* web tool ^50^. *CircInteractome* uses TargetScan’s Perl script for prediction of microRNA with exact 7-mer or 8-mer seed sequence, complementing the mature circular RNA sequence. Cytoscape was used to visualize circular RNA-microRNA interaction. For circular RNAs acting as microRNA precursor, the mature sequences of circular RNA were searched for stem-loop microRNA sequence using BLAST tool in mirBASE.

## RESULTS

### Expression signature of circular RNAs in breast cancer

RNA sequencing of breast tissues resulted in approximately 89 million reads which were filtered for unmapped reads using bowtie2. The unmapped reads were used for identification of circular RNAs using two different algorithms, find_circ and DCC. Find_Circ tool was used to identify circular RNAs by converting unmapped reads to the anchor fastq file using unmapped2anchors.py. The identified number of circular RNAs varied between samples across all groups with a minimum number of 2958 circular RNAs to a maximum of 21214 circular RNAs. About 1769 circular RNAs were found to carry unique back spliced reads in at least three samples and circular RNAs left after filtering by reads in individual sample ranges from 560 to 1632. Likewise, 1413 overlapping circular RNAs were detected by DCC analysis, which ranges from 1179 to 1412 among the samples [Table1].

The genomic features of identified circular RNAs show that exonic circular RNAs are the most abundant class accounting for 78 % obtained from find_circ and 75.8 % from DCC. Circular RNAs consisting of 5’UTR-exon (13.1% & 14.6%) and exon-3’UTR sequence (2% & 2.7%) also contributed to the number of identified circular RNAs through find_circ and DCC respectively [Figure 1]. Circular RNAs arising from exon-intron and intergenic regions and others (5’UTR-3’UTR, UTR-tx, exon-tx etc) were ranging from 2.9 – 0.5%. The results show that majority of circular RNAs carry exon-exon sequence arising due to mRNA splicing and circularization.

**Figure 1:**
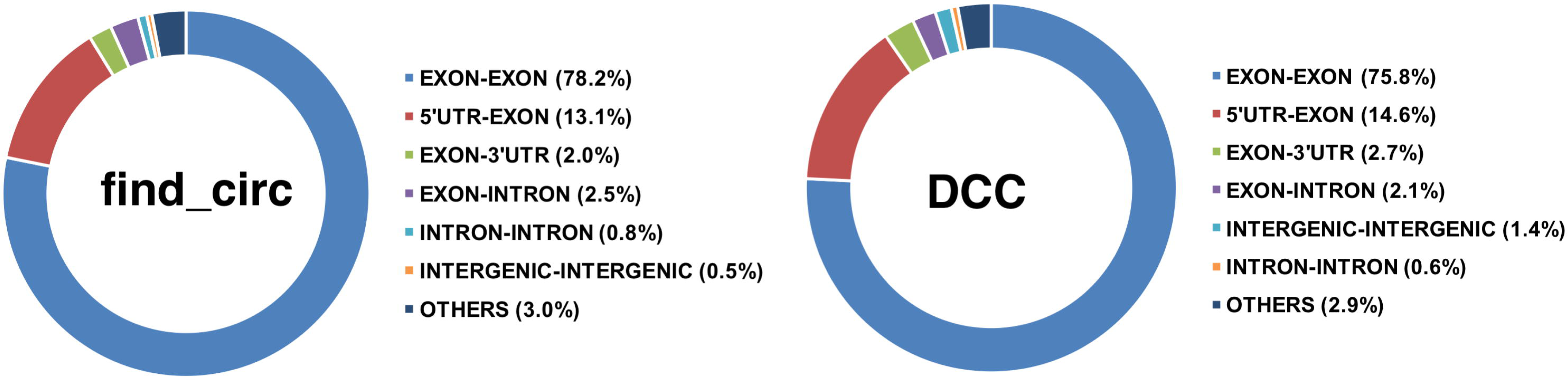
Genomic features of identified circular RNAs in early stage breast cancer

### Differentially expressed circular RNAs in breast tumors

Among 1769 and 1413 circular RNAs identified using find_circ and DCC, respectively, 1516 and 1270 circular RNAs were annotated using CircBase reference. Hierarchal clustering shows the grouping of samples distinctively into tumor-matched normal and normal illustrating the characteristic expression of circular RNAs associated with breast cancer [Figure 2]. Almost all of the down-regulated circular RNAs (hsa_circ_0001064, hsa_circ_0001073, hsa_circ_0001097, hsa_circ_0001400 etc.) were expressed at relatively low levels in tumor and matched normal tissues in comparison to normal tissues. However, two circular RNAs, hsa_circ_7386 (circCRIM1) and hsa_circ_8345 (circPTK2) were downregulated in tumor relative to both matched normal and normal tissue samples. The results establish breast cancer specific circular RNAs deregulated in tumor compared to normal tissue obtained from the same patient. Expression variation between normal breast tissue and tumor tissue illustrate distinct circular RNA signature profile specific to early stage breast cancer. DEseq2 analysis on circular RNA identified by find_circ showed 58 differentially regulated circular RNAs (30 upregulated and 28 downregulated), whereas 87 deregulated circular RNAs consisting of 61 upregulated and 26 downregulated candidates resulted from DCC-identified circular RNAs. Several common deregulated circular RNAs were found when tumor samples were compared to matched normal and normal separately (Supplementary Table 1 and 2). In total 26 circular RNAs (13 upregulated and 13 downregulated) were found to be differentially regulated in both find_circ and DCC based analysis [Table 2].

**Table 1:**
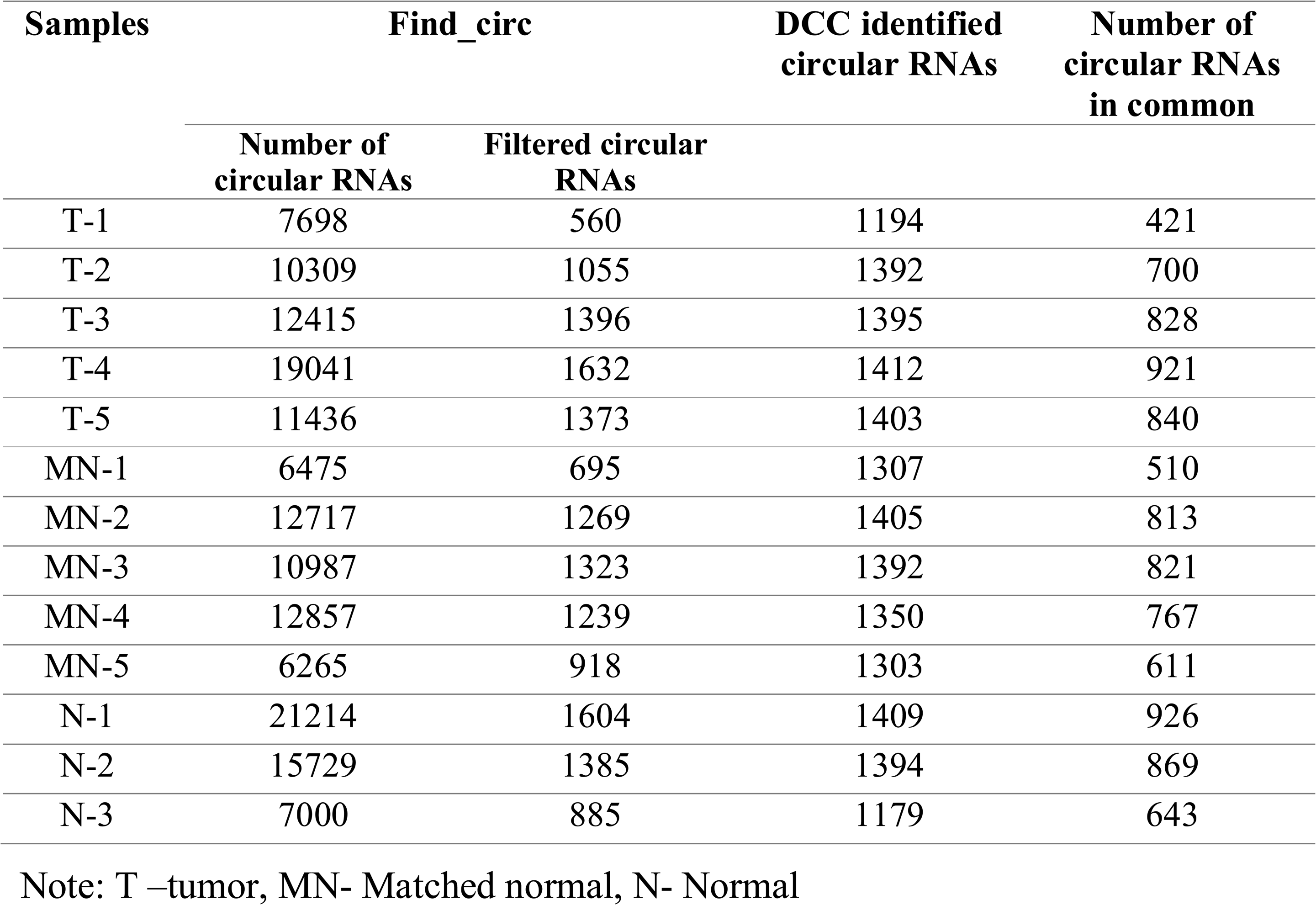
Identified number of circular RNAs

**Table 2.** List of differentially regulated circular RNAs in early stage breast cancer

| Circular RNA | MN-T vs N<br>(log<sub>2</sub>FC) |  | T vs MN<br>(log<sub>2</sub>FC) |  | T vs N<br>(log<sub>2</sub>FC) |  | Parental Gene |
| --- | --- | --- | --- | --- | --- | --- | --- |
|  | DCC | find_circ | DCC | find_circ | DCC | find_circ |  |
| hsa_circ_0000117 | 1.13 | 1.08 | - | - | 1.11 | 1.05 | MAN1A2 |
| hsa_circ_0000283 | 1.65 | 2.63 | - | - | - | - | CSTF3 |
| hsa_circ_0000639 | - | - | - | - | 2.05 | 2.61 | ETFA |
| hsa_circ_0001064 | -1.17 | -2.68 | - | - | - | - | EPB41L5 |
| hsa_circ_0001073 | - | - | - | - | -1.56 | -1.32 | ACVR2A |
| hsa_circ_0001097 | -1.37 | -1.51 | - | - | - | - | PIKFYVE |
| hsa_circ_0001400 | -1.25 | -1.35 | - | - | -1.20 | -1.36 | RELL1 |
| hsa_circ_0001519 | -1.24 | -1.45 | - | - | -1.25 | -1.34 | MAN2A1 |
| hsa_circ_0001946 | 2.10 | 1.58 | - | - | 2.00 | 1.47 | AL078639.1 |
| hsa_circ_0002346 | - | - | - | - | -2.10 | -1.85 | CRIM1 |
| hsa_circ_0002496 | 2.28 | 4.02 | - | - | 2.42 | 4.10 | APPBP1 |
| hsa_circ_0003247 | - | - | - | - | 1.19 | 1.09 | ASH1L |
| hsa_circ_0004069 | -1.30 | -1.43 | - | - | -1.72 | -2.01 | SLC37A3 |
| hsa_circ_0006743 | 2.18 | 4.77 | - | - | - | - | JMJD1C |
| hsa_circ_0006884 | -1.17 | -1.5 | - | - | -1.12 | -1.45 | TIMMDC1 |
| hsa_circ_0006950 | - | - | 1.37 | 1.97 | - | - | TRPS1 |
| hsa_circ_0007386 | - | - | - | - | -1.71 | -1.65 | CRIM1 |
| hsa_circ_0007766 | - | - | 3.14 | 3.03 | - | - | ERBB2 |
| hsa_circ_0007897 | -1.17 | -1.98 | - | - | - | - | BOC |
| hsa_circ_0008225 | - | - | 2.19 | 2.49 | 1.96 | 2.01 | ZMYND11 |
| hsa_circ_0008311 | 1.13 | 1.25 | - | - | - | - | STAM |
| hsa_circ_0008345 | - | - | - | - | -1.29 | -1.64 | PTK2 |
| hsa_circ_0009069 | -1.06 | -1.61 | - | - | - | - | PHF8 |
| hsa_circ_0016601 | - | - | 3.58 | 3.58 | - | - | DNAH14 |
| hsa_circ_0023990 | 1.72 | 2.43 | - | - | 1.99 | 2.85 | NOX4 |
| hsa_circ_0025765 | -1.59 | -1.73 | - | - | -2.01 | -1.96 | TMTC1 |

Note: T –tumor, MN- Matched normal, N- Normal; Log2FC - p< 0.05

**Table 3.**
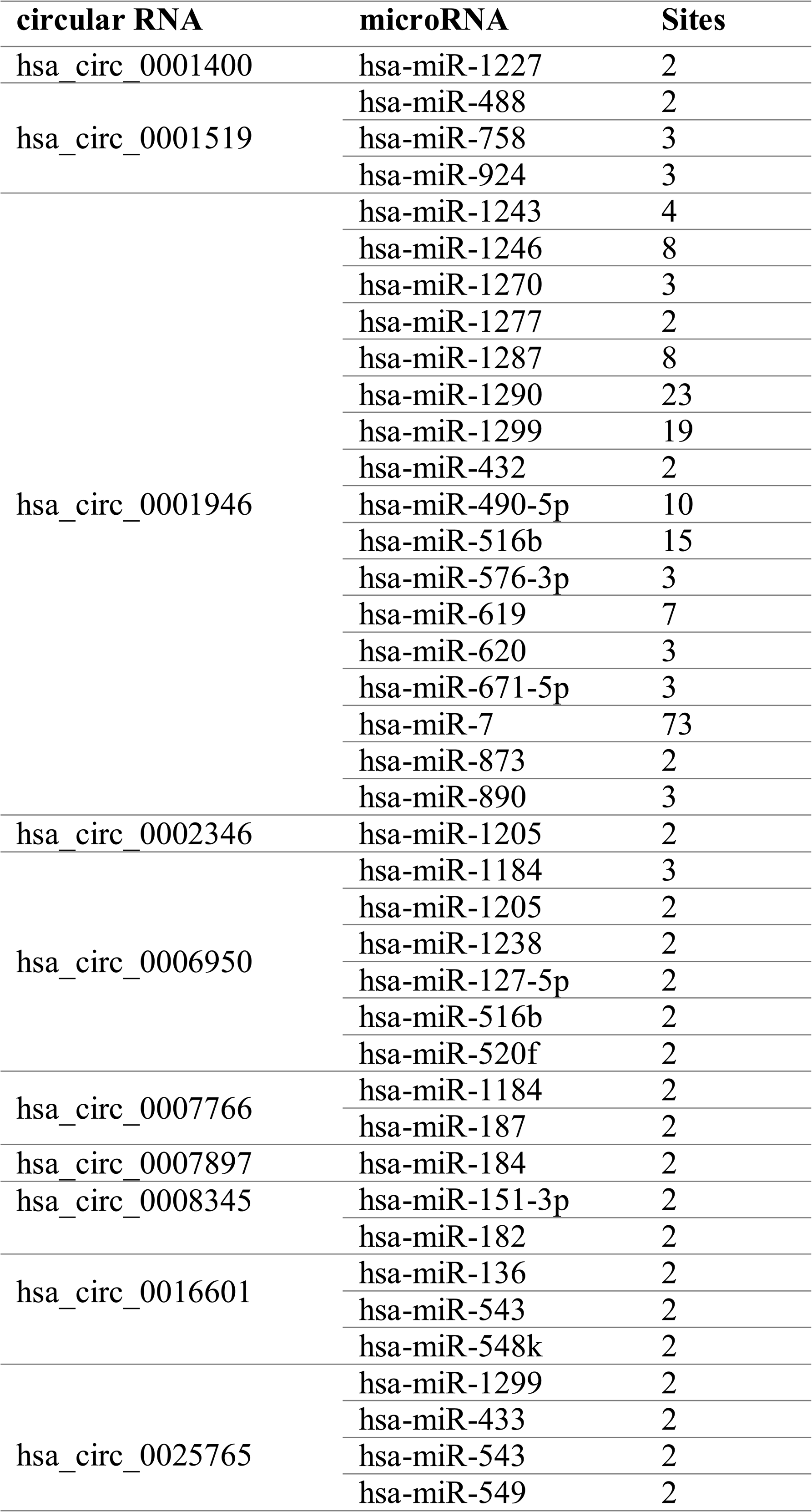
List of circular RNAs with predicted number of miRNA binding sites

**Figure 2:**
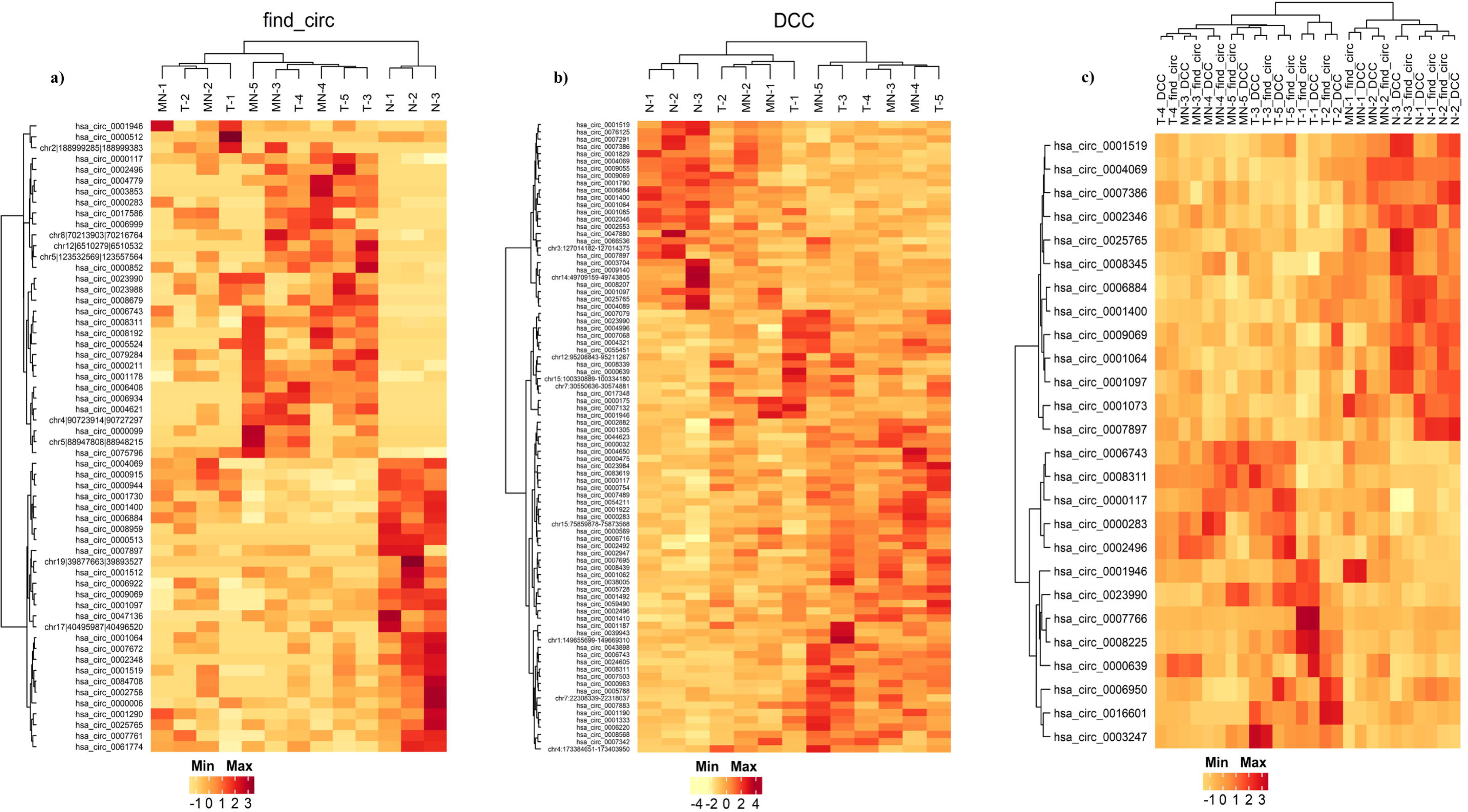
a) Heatmap of differentially expressed circular RNAs by find_circ (N= 58) b) Heatmap of differentially expressed circular RNAs by DCC tool (N= 87), c) commonly deregulated circular RNAs in both find_circ and DCC in early stage breast cancer (N= 26)

Based on normalized read counts from both pipelines, the expression status was plotted to specifically identify candidate circular RNAs differentially expressed in early stage breast cancer. Among the upregulated circular RNAs, hsa_circ_0001946 (*CDR1as*) showed a high expression in tumor tissues and matched normal tissue when compared with normal. Similarly, hsa_circ_0000117 (circMAN1A2) and hsa_circ_0023990 (circNOX4) were also observed with increased expression levels in both tumor and matched normal tissue. Hsa_circ_0008225 (circZMYND11) and hsa_circ_0016601 (circDNAH14) were specifically upregulated in tumor tissues compared to matched normal and normal tissue.

### MicroRNA sponging by circular RNA

The sponging potential of circular RNA was predicted using the web tool *CircInteractome.* Exact seed sequence match of 7mers or 8mers for the 26 circular RNAs that were differentially expressed in both find_circ and DCC were predicted. About 223 microRNA were predicted to be sponged by these 26 circular RNAs through 495 interactions. About 40 interactions were filtered by considering circular RNAs carrying at least 2 binding sites for single microRNA [Figure 3]. Hsa_circ_0001946 was found to harbor the highest number of microRNA binding sites with 73 binding regions for miR-7. We speculate that hsa_circ_0001946 may be a specific biomarker for breast cancer with possible functional role through sponging of microRNA. In addition, hsa_circ_0006950 was predicted with 2-3 binding sites for about 6 microRNA. Similarly, hsa_circ_00256765 showed binding sites for four microRNA; hsa_circ_0016001 and hsa_circ_0001519 harbored binding sites for 3 microRNA each [Table 3]. Hence we hypothesized that the identified differentially expressed circular RNAs carrying multiple microRNA binding sites could alter microRNA function by sponging.

**Figure 3:**
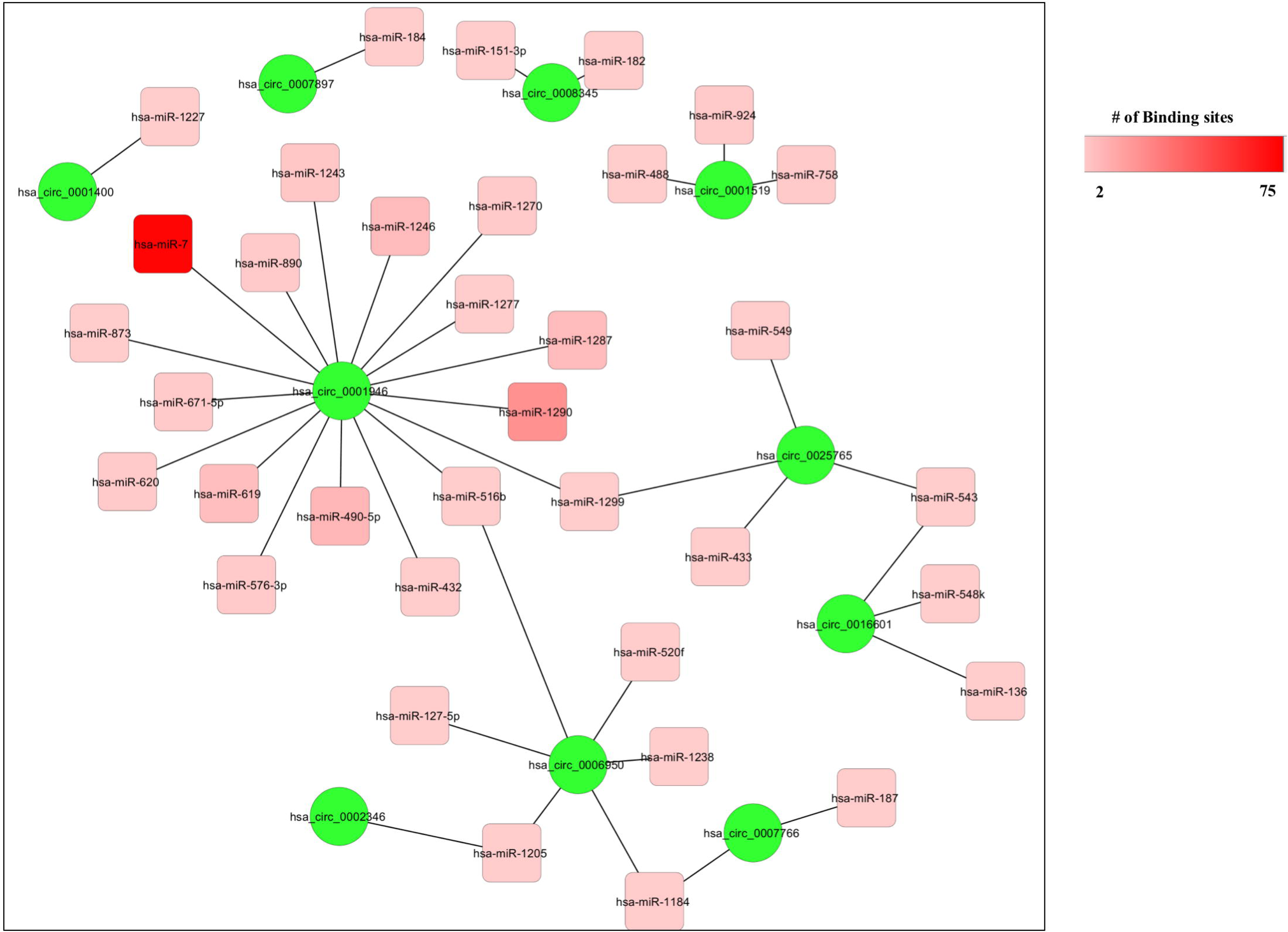
Network of circular RNAs and binding microRNAs in early stage breast cancer. Green circles represent circular RNAs and round squares represent microRNAs, intensity marked by colour gradation indicate microRNA binding sites.

### Circular RNA act as precursors for microRNA

The secondary structure of circular RNAs showed several stem loop structures resembling pre-microRNA structure. The mature circular RNA sequences were checked for carrying pre-microRNA that can generate microRNA. Of the circular RNAs identified, hsa_circ_0001519 was found to contain precursor sequences for 5 microRNA while hsa_circ_0007766 contained precursor sequences for 4 microRNA [Table 4]. Five circular RNAs, namely hsa_circ_0002346, hsa_circ_0001064, hsa_circ_0002496, hsa_circ_0000639 and hsa_circ_0001073, were predicted to yield 3 microRNA each. Hsa_circ_0007766 was predicted to give rise to miR-370, commonly upregulated microRNA in breast cancer. Besides, miR-141 and miR-372 were predicted to be processed from circ_0008225 and circ_0002496 respectively had been previously reported to possess tumor suppressive property.

**Table 4.** List of circular RNAs predicted to be a precursor for microRNAs

| **Circular RNA** | **Number of microRNAs**<br>**originated** | **MicroRNA** |
| --- | --- | --- |
| hsa_circ_0007766 | 3 | hsa-miR-6865; hsa-miR-3155a; hsa-miR-370 |
| hsa_circ_0002346 | 3 | hsa-miR-1263; hsa-miR-12118 ; hsa-miR-7114 |
| hsa_circ_0008225 | 2 | hsa-miR-141; hsa-miR-19a |
| hsa_circ_0001064 | 3 | hsa-miR-3612; hsa-miR-4447; hsa-miR-9986 |
| hsa_circ_0008311 | 2 | hsa-miR-6504; hsa-miR-517a |
| hsa_circ_0001097 | 2 | hsa-miR-211; hsa-miR-548ay |
| hsa_circ_0001519 | 5 | hsa-miR-5195; hsa-miR-3913-1; hsa-miR-3913-2 ; hsa-miR-6864; hsa-miR-379 |
| hsa_circ_0002496 | 3 | hsa-miR-372; hsa-miR-450b; hsa-miR-450a-2 |
| hsa_circ_0006884 | 2 | hsa-miR-3152; hsa-miR-548am |
| hsa_circ_0007897 | 2 | hsa-miR-1266; hsa-miR-3085 |
| hsa_circ_0000639 | 3 | hsa-miR-2681; hsa-miR-548k ; hsa-miR-4299 |
| hsa_circ_0001073 | 3 | hsa-miR-6129; hsa-miR-449a; hsa-miR-563 |
| hsa_circ_0007386 | 2 | hsa-miR-1263; hsa-miR-7114 |

## DISCUSSION

Circular RNAs are emerging non-coding RNA recognized with immense potential in controlling regulatory events, playing significant role in different diseases including cancer. Several studies have established the differential expression of circular RNAs in tumor tissues compared to normal tissue in a range of cancer types ^23^. Circular RNAs were also detectable in circulation especially enriched in exosomes indicating their significance as non-invasive tumor biomarkers ^16,24^. Recent findings on circular RNA/microRNA axis of gene expression regulation demonstrate higher degree of complexity. Therefore, interrogating circular RNAs can uncover multiple fundamental mechanisms in cancer. Identification of circular RNAs is challenging with the current tools available. Although we have several pipelines for identifying circular RNAs, all of them are back-spliced junction based detection methods with minor variations ^25^.

In our study, we combined the power of RNA sequencing that enables a comprehensive analysis of RNA expression across the transcriptome, with two different pipelines to detect circular RNAs in a set of tumor, matched normal and normal samples. Find_circ utilizes the unmapped reads from bowtie2 to identify back-spliced junctions while DCC detects circular RNA through splice aware aligner STAR. The number of deregulated circular RNAs varied between find_circ and DCC but 944 common circular RNAs were found among a total of 1769 (find_circ) and 1413 (DCC) predicted circular RNAs. It was interesting to observe that the genomic landscape of most circular RNAs was exonic followed by 5’UTR-exon and exon-3’UTR, implicating major contribution from mRNA splicing. The circular RNA transcripts with intron and UTR sequence can sequester microRNA especially the exonic-3’UTR circular RNAs which may harbor multiple microRNA response elements. Upregulation of such circular RNAs will compete for microRNA affecting target mRNA levels. In this study, early stage breast tumors (Stage I-IIA) were taken for investigation along with matched normal and normal tissues to delineate relevant circular RNAs. Most importantly, differentially regulated circular RNAs among tumor-matched normal samples indicated prime circular RNA candidates (hsa_circ_0002496 and hsa_circ_0006743) that are deregulated in individuals with cancer or having a predisposition for the disease. These circular RNAs, apart from diagnostic use, can be verified for risk association with breast cancer. Similarly, hsa_circ_0000639 and hsa_circ_0025765 are novel circular RNAs identified to be upregulated in tumor tissues compared to normal breast tissue. Certain circular RNAs (hsa_circ_0007766 and hsa_circ_0016601) were overexpressed exclusively in tumor tissues when compared to the adjacent normal tissue. However, these circular RNAs when associated with subtypes can be more informative on treatment response. Recently, circular RNA expression was evaluated using TCGA-BRCA data correlating subtypes and risk of relapse, indicative on the use of circular RNAs in clinical application ^26^. Moreover, few circular RNAs like circRNA-MTO1 and circ_0006528 have been associated with chemo-resistance in breast cancer ^27,28^.

Circular RNA-microRNA interaction has proved to be a potential mechanism of gene regulation that plays significant role in breast cancer progression. The sponging of mir-1271 and miR-143 by circ-ABCB10 and hsa_circ_0001982 respectively has been demonstrated earlier with breast cancer ^21,22^. In our analysis, we observed that hsa_circ_0001946, also known as CDR1as, is differentially regulated in early stage breast tumors. Overexpression of CDR1as decoys miR-7 with its 73 binding sites causing tumor promotion through epithelial-to-mesenchymal transition, invasion and metastasis ^29-31^. Oncogenic potential of CDR1as through miR-7 sponging has been shown in gastric cancer and carcinoma of liver, esophagus and lung ^19,32-34^. Since miR-7 has several proven targets like CCNE1, PIK3CD and KLF4, one or more pathways seems to be affected by CDR1as. Besides miR-7, CDR1as also sponges other microRNAs like miR-135a, miR-1290, miR-876-5p, miR-671-5p ^35-37^. Nevertheless, dysregulated CDR1as expression and sponging of particular microRNA hugely determine their functional role in tumor development. Similarly, hsa_circ_0006950 was another circular RNA overexpressed in our study samples that was predicted with 6 microRNA binding sites. Notably, microRNA like miR-1184, miR-127-5p and miR-516b that can be sponged by the identified circular RNAs have previously been associated with prognosis of breast cancer ^38-40^. Other circular RNAs interacting microRNA such as miR-433, miR-151, miR-182, miR-136 and miR-543 have been previously associated with breast cancer development ^41-45^. Under normal physiological condition, most of the circular RNAs are expressed at low levels ^46^. However, high expression of circular RNAs can determine microRNA activity by either decoying or utilizing Argonaute-associated microRNA forming the silencing complex. The expression of circular RNAs can hence be detrimental to the expression or availability of such microRNA. Nevertheless, it is warranted to evaluate the regulatory relationship between circular RNA and microRNA experimentally. Although we have identified novel circular RNAs differentially expressed in early stage breast cancer, the main limitation of our study is the requirement of validation in a larger series of samples with clinical correlation. Another major limitation of the study is adapting the mature circular RNA sequence retrieved from database instead of the sequence read from our analysis. As a result, any novel splice junction creating new microRNA binding sites from our analysis may not be addressed.

In summary, we have used two different circular RNA detection tools to identify aberrantly expressed circular RNAs in breast cancer. We have identified 26 circular RNAs differentially expressed between tumor and normal. Further, we have explored circular RNA-microRNA interaction and reported multiple microRNA binding sites that can aid in its sponging function. Interestingly, some circular RNAs were found to harbor pre-microRNA sequence, thus having possibility to act as precursors for microRNA of relevance in breast cancer. This study identifies novel breast cancer associated circular RNAs that can be explored for diagnostic and prognostic potential.

## Author Contributions

Conceptualization, S.M and A.K.D.M.R; Methodology, V.R.A., B.M., V.S., S.S. and S.M; Software, V.R.A; Validation, A.K.D.M.R, V.R.A and S.M.; Formal Analysis, A.K.D.M.R and V.R.A.; Investigation, S.M.; Resources, V.S. and S.S.; Data Curation, P.R.; Writing – Original Draft Preparation, A.K.D.M.R; Writing – Review & Editing, P.R., S.M., T.R. and Z.H.; Visualization, V.R.A. and A.K.D.M.R.; Supervision, T.R. and S.M.; Project Administration, B.M. and S.M.; Funding Acquisition, S.M.

## Supplementary Materials

Supplementary Table 1: List of differentially regulated circRNA identified by find_circ between tumor, matched normal samples and normal samples. Supplementary Table 2: List of differentially regulated circRNA identified by DCC between tumor, matched normal samples and normal samples. Supplementary Table 3. Clinico-pathological characteristics of samples

## References

1. Ferlay J SI, Ervik M, Dikshit R, Eser S, Mathers C, Rebelo M, Parkin DM, Forman D, Bray, F. GLOBOCAN 2012 v1.0, Cancer Incidence and Mortality Worldwide: IARC CancerBase No. 11 [Internet]. Lyon, France: International Agency for Research on Cancer; 2013 Available from: http://globocaniarcfr/,. accessed on 06/08/2018.

2. Anastasiadou E, Jacob LS, Slack FJ. Non-coding RNA networks in cancer. Nat Rev Cancer. 2018;18(1):5–18.

3. Bhan A, Soleimani M, Mandal SS. Long Noncoding RNA and Cancer: A New Paradigm. Cancer Res. 2017;77(15):3965–3981.

4. Hayes J, Peruzzi PP, Lawler S. MicroRNAs in cancer: biomarkers, functions and therapy. Trends Mol Med. 2014;20(8):460–469.

5. Iorio MV, Croce CM. MicroRNA dysregulation in cancer: diagnostics, monitoring and therapeutics. A comprehensive review. EMBO Mol Med. 2012;4(3):143–159.

6. Danan M, Schwartz S, Edelheit S, Sorek R. Transcriptome-wide discovery of circular RNAs in Archaea. Nucleic Acids Res. 2012;40(7):3131–3142.

7. Wang Y, Wang Z. Efficient backsplicing produces translatable circular mRNAs. Rna. 2015;21(2):172–179.

8. Chen X, Han P, Zhou T, Guo X, Song X, Li Y. circRNADb: A comprehensive database for human circular RNAs with protein-coding annotations. Sci Rep. 2016;6:34985.

9. Glazar P, Papavasileiou P, Rajewsky N. circBase: a database for circular RNAs. Rna. 2014;20(11):1666–1670.

10. Hou LD, Zhang J. Circular RNAs: An emerging type of RNA in cancer. Int J Immunopathol Pharmacol. 2017;30(1):1–6.

11. Xu H, Gong Z, Shen Y, Fang Y, Zhong S. Circular RNA expression in extracellular vesicles isolated from serum of patients with endometrial cancer. Epigenomics. 2018;10(2):187–197.

12. Coscujuela Tarrero L, Ferrero G, Miano V, et al. Luminal breast cancer-specific circular RNAs uncovered by a novel tool for data analysis. Oncotarget. 2018;9(18):14580–14596.

13. Du WW, Fang L, Yang W, et al. Induction of tumor apoptosis through a circular RNA enhancing Foxo3 activity. Cell Death Differ. 2017;24(2):357–370.

14. Lu L, Sun J, Shi P, et al. Identification of circular RNAs as a promising new class of diagnostic biomarkers for human breast cancer. Oncotarget. 2017;8(27):44096–44107.

15. Bahn JH, Zhang Q, Li F, et al. The landscape of microRNA, Piwi-interacting RNA, and circular RNA in human saliva. Clin Chem. 2015;61(1):221–230.

16. Li Y, Zheng Q, Bao C, et al. Circular RNA is enriched and stable in exosomes: a promising biomarker for cancer diagnosis. Cell Res. 2015;25(8):981–984.

17. Lasda E, Parker R. Circular RNAs: diversity of form and function. RNA. 2014;20(12):1829–1842.

18. Hansen TB, Wiklund ED, Bramsen JB, et al. miRNA-dependent gene silencing involving Ago2-mediated cleavage of a circular antisense RNA. Embo j. 2011;30(21):4414–4422.

19. Yu L, Gong X, Sun L, Zhou Q, Lu B, Zhu L. The Circular RNA Cdr1as Act as an Oncogene in Hepatocellular Carcinoma through Targeting miR-7 Expression. PLoS One. 2016;11(7):e0158347.

20. Wu J, Jiang Z, Chen C, et al. CircIRAK3 sponges miR-3607 to facilitate breast cancer metastasis. Cancer Lett. 2018;430:179–192.

21. Tang YY, Zhao P, Zou TN, et al. Circular RNA hsa_circ_0001982 Promotes Breast Cancer Cell Carcinogenesis Through Decreasing miR-143. DNA Cell Biol. 2017;36(11):901–908.

22. Liang HF, Zhang XZ, Liu BG, Jia GT, Li WL. Circular RNA circ-ABCB10 promotes breast cancer proliferation and progression through sponging miR-1271. Am J Cancer Res. 2017;7(7):1566–1576.

23. Geng Y, Jiang J, Wu C. Function and clinical significance of circRNAs in solid tumors. J Hematol Oncol. 2018;11(1):98.

24. Yin WB, Yan MG, Fang X, Guo JJ, Xiong W, Zhang RP. Circulating circular RNA hsa_circ_0001785 acts as a diagnostic biomarker for breast cancer detection. Clin Chim Acta. 2017.

25. Gao Y, Zhao F. Computational Strategies for Exploring Circular RNAs. Trends Genet. 2018;34(5):389–400.

26. Nair AA, Niu N, Tang X, et al. Circular RNAs and their associations with breast cancer subtypes. Oncotarget. 2016;7(49):80967–80979.

27. Gao D, Zhang X, Liu B, et al. Screening circular RNA related to chemotherapeutic resistance in breast cancer. Epigenomics. 2017;9(9):1175–1188.

28. Liu Y, Dong Y, Zhao L, Su L, Luo J. Circular RNAMTO1 suppresses breast cancer cell viability and reverses monastrol resistance through regulating the TRAF4/Eg5 axis. Int J Oncol. 2018.

29. Hansen TB, Jensen TI, Clausen BH, et al. Natural RNA circles function as efficient microRNA sponges. Nature. 2013;495(7441):384–388.

30. Kong X, Li G, Yuan Y, et al. MicroRNA-7 inhibits epithelial-to-mesenchymal transition and metastasis of breast cancer cells via targeting FAK expression. PLoS One. 2012;7(8):e41523.

31. Yu N, Huangyang P, Yang X, et al. microRNA-7 suppresses the invasive potential of breast cancer cells and sensitizes cells to DNA damages by targeting histone methyltransferase SET8. J Biol Chem. 2013;288(27):19633–19642.

32. Pan H, Li T, Jiang Y, et al. Overexpression of Circular RNA ciRS-7 Abrogates the Tumor Suppressive Effect of miR-7 on Gastric Cancer via PTEN/PI3K/AKT Signaling Pathway. J Cell Biochem. 2018;119(1):440–446.

33. Zhang X, Yang D, Wei Y. Overexpressed CDR1as functions as an oncogene to promote the tumor progression via miR-7 in non-small-cell lung cancer. Onco Targets Ther. 2018;11:3979–3987.

34. Huang H, Wei L, Qin T, Yang N, Li Z, Xu Z. Circular RNA ciRS-7 triggers the migration and invasion of esophageal squamous cell carcinoma via miR-7/KLF4 and NF-kappaB signals. Cancer Biol Ther. 2018:1–8.

35. Li P, Yang X, Yuan W, et al. CircRNA-Cdr1as Exerts Anti-Oncogenic Functions in Bladder Cancer by Sponging MicroRNA-135a. Cell Physiol Biochem. 2018;46(4):1606– 1616.

36. Sang M, Meng L, Sang Y, et al. Circular RNA ciRS-7 accelerates ESCC progression through acting as a miR-876-5p sponge to enhance MAGE-A family expression. Cancer Lett. 2018;426:37–46.

37. Barbagallo D, Condorelli A, Ragusa M, et al. Dysregulated miR-671-5p / CDR1-AS / CDR1 / VSNL1 axis is involved in glioblastoma multiforme. Oncotarget. 2016;7(4):4746– 4759.

38. Tahiri A, Leivonen SK, Luders T, et al. Deregulation of cancer-related miRNAs is a common event in both benign and malignant human breast tumors. Carcinogenesis. 2014;35(1):76–85.

39. Farina NH, Ramsey JE, Cuke ME, et al. Development of a predictive miRNA signature for breast cancer risk among high-risk women. Oncotarget. 2017;8(68):112170– 112183.

40. Wang S, Li H, Wang J, Wang D, Yao A, Li Q. Prognostic and biological significance of microRNA-127 expression in human breast cancer. Dis Markers. 2014;2014:401986.

41. Zhang T, Jiang K, Zhu X, et al. miR-433 inhibits breast cancer cell growth via the MAPK signaling pathway by targeting Rap1a. Int J Biol Sci. 2018;14(6):622–632.

42. Yeh TC, Huang TT, Yeh TS, et al. miR-151-3p Targets TWIST1 to Repress Migration of Human Breast Cancer Cells. PLoS One. 2016;11(12):e0168171.

43. Zhang X, Ma G, Liu J, Zhang Y. MicroRNA-182 promotes proliferation and metastasis by targeting FOXF2 in triple-negative breast cancer. Oncol Lett. 2017;14(4):4805–4811.

44. Yan M, Li X, Tong D, et al. miR-136 suppresses tumor invasion and metastasis by targeting RASAL2 in triple-negative breast cancer. Oncol Rep. 2016;36(1):65–71.

45. Chen P, Xu W, Luo Y, et al. MicroRNA 543 suppresses breast cancer cell proliferation, blocks cell cycle and induces cell apoptosis via direct targeting of ERK/MAPK. Onco Targets Ther. 2017;10:1423–1431.

46. Memczak S, Jens M, Elefsinioti A, et al. Circular RNAs are a large class of animal RNAs with regulatory potency. Nature. 2013;495(7441):333–338.

47. Legnini I, Di Timoteo G, Rossi F, et al. Circ-ZNF609 Is a Circular RNA that Can Be Translated and Functions in Myogenesis. Mol Cell. 2017;66(1):22–37 e29.

48. Cheng J, Metge F, Dieterich C. Specific identification and quantification of circular RNAs from sequencing data. Bioinformatics. 2016;32(7):1094–1096.

49. Dobin A, Davis CA, Schlesinger F, et al. STAR: ultrafast universal RNA-seq aligner. Bioinformatics. 2013;29(1):15–21.

50. Dudekula DB, Panda AC, Grammatikakis I, De S, Abdelmohsen K, Gorospe M. CircInteractome: A web tool for exploring circular RNAs and their interacting proteins and microRNAs. RNA Biol. 2016;13(1):34–42.

