## Supplementary Table 1 and 2 for "RNA sequencing identifies dysregulated circular RNAs in early-stage breast cancer"

1. List of differentially regulated circRNA identified by find_circ - tumor -matched normal samples compared to normal samples

| CircRNA | baseMean | log2FoldChange | pvalue |
|---|---|---|---|
| hsa_circ_0000099 | 1.845 | 3.955 | 0.0323646 |
| hsa_circ_0000117 | 15.831 | 1.082 | 0.0048419 |
| hsa_circ_0000006 | 5.165 | -2.008 | 0.0183003 |
| hsa_circ_0008311 | 7.731 | 1.250 | 0.0401526 |
| hsa_circ_0006408 | 1.359 | 3.590 | 0.0283160 |
| hsa_circ_0008679 | 3.065 | 4.696 | 0.0019965 |
| hsa_circ_0017586 | 1.173 | 3.399 | 0.0368795 |
| hsa_circ_0006743 | 3.246 | 4.765 | 0.0005649 |
| hsa_circ_0002758 | 1.631 | -2.231 | 0.0445063 |
| hsa_circ_0000211 | 28.682 | 1.063 | 0.0073984 |
| hsa_circ_0000283 | 2.242 | 2.626 | 0.0436868 |
| hsa_circ_0023988 | 3.115 | 4.679 | 0.0011460 |
| hsa_circ_0023990 | 4.079 | 2.429 | 0.0394845 |
| hsa_circ_0025765 | 3.438 | -1.732 | 0.0196106 |
| chr12\|6510279\|6510532 | 1.875 | 4.021 | 0.0277914 |
| hsa_circ_0000512 | 2.561 | -3.386 | 0.0053897 |
| hsa_circ_0000513 | 1.030 | -4.599 | 0.0037716 |
| chr17\|40495987\|40496520 | 1.544 | -2.569 | 0.0447307 |
| hsa_circ_0004621 | 1.623 | 3.824 | 0.0178141 |
| hsa_circ_0002496 | 1.824 | 4.016 | 0.0123018 |
| hsa_circ_0047136 | 1.852 | -2.025 | 0.0273281 |
| hsa_circ_0000852 | 4.280 | 1.995 | 0.0353462 |
| hsa_circ_0000915 | 18.404 | -1.239 | 0.0247590 |
| chr19\|39877663\|39893527 | 7.240 | -7.407 | 0.0000896 |
| hsa_circ_0000944 | 10.671 | -1.188 | 0.0214889 |
| hsa_circ_0001064 | 2.051 | -2.683 | 0.0077155 |
| chr2\|188999285\|188999383 | 2.223 | 4.163 | 0.0291970 |
| hsa_circ_0001097 | 4.017 | -1.506 | 0.0166777 |
| hsa_circ_0006934 | 1.512 | 3.751 | 0.0203724 |
| hsa_circ_0008959 | 1.100 | -3.847 | 0.0080867 |
| hsa_circ_0002348 | 0.864 | -3.503 | 0.0271637 |
| hsa_circ_0003853 | 1.309 | 3.529 | 0.0494910 |
| hsa_circ_0006922 | 4.305 | -1.161 | 0.0431035 |
| hsa_circ_0001178 | 15.056 | 1.028 | 0.0057221 |
| hsa_circ_0061774 | 5.364 | -1.173 | 0.0494319 |
| hsa_circ_0007897 | 3.058 | -1.980 | 0.0398100 |
| hsa_circ_0006884 | 5.257 | -1.496 | 0.0067902 |
| hsa_circ_0006999 | 1.589 | 3.824 | 0.0178555 |
| hsa_circ_0001290 | 6.082 | -1.286 | 0.0222771 |

| | | | |
|---|---|---|---|
| hsa_circ_0007761 | 5.638 | -1.341 | 0.0315121 |
| hsa_circ_0001400 | 16.690 | -1.352 | 0.0006869 |
| chr4\|90723914\|90727297 | 1.426 | 3.653 | 0.0266736 |
| hsa_circ_0001519 | 8.444 | -1.454 | 0.0265411 |
| chr5\|123532569\|123557564 | 1.413 | 3.683 | 0.0196608 |
| hsa_circ_0005524 | 1.708 | 3.783 | 0.0294885 |
| chr5\|88947808\|88948215 | 1.519 | 3.690 | 0.0481955 |
| hsa_circ_0001512 | 1.479 | -2.814 | 0.0268953 |
| hsa_circ_0075796 | 13.540 | 1.044 | 0.0061488 |
| hsa_circ_0001730 | 10.605 | -1.433 | 0.0021331 |
| hsa_circ_0004069 | 7.548 | -1.425 | 0.0390599 |
| hsa_circ_0004779 | 2.265 | 4.299 | 0.0085096 |
| hsa_circ_0079284 | 1.771 | 3.916 | 0.0118362 |
| hsa_circ_0007672 | 2.402 | -2.242 | 0.0417748 |
| hsa_circ_0084708 | 1.696 | -2.443 | 0.0461953 |
| chr8\|70213903\|70216764 | 13.331 | 1.113 | 0.0124415 |
| hsa_circ_0008192 | 1.875 | 3.985 | 0.0305754 |
| hsa_circ_0001946 | 70.775 | 1.583 | 0.0253275 |
| hsa_circ_0009069 | 3.451 | -1.606 | 0.0146519 |

2. List of differentially regulated circRNA identified by find_circ - tumor samples compared to matched normal samples

| CircRNA | baseMean | log2FoldChange | pvalue |
|---|---|---|---|
| hsa_circ_0000110 | 2.137 | 2.778 | 0.0443544 |
| hsa_circ_0016601 | 3.855 | 3.585 | 0.0004574 |
| hsa_circ_0005899 | 1.047 | 3.436 | 0.0407840 |
| chr10\|11851997\|11852215 | 1.735 | -3.453 | 0.0228011 |
| hsa_circ_0008225 | 4.062 | 2.488 | 0.0036967 |
| chr11\|119296918\|119297481 | 1.171 | 3.514 | 0.0331122 |
| hsa_circ_0007401 | 1.247 | 3.191 | 0.0281106 |
| chr12\|28255581\|28307749 | 1.461 | 3.201 | 0.0248117 |
| hsa_circ_0008776 | 4.445 | -1.651 | 0.0313786 |
| hsa_circ_0033475 | 1.597 | -3.914 | 0.0097891 |
| hsa_circ_0008603 | 1.355 | -4.113 | 0.0232940 |
| hsa_circ_0007766 | 7.818 | 3.031 | 0.0052298 |
| chr17\|40495987\|40496520 | 1.544 | -3.183 | 0.0206316 |
| hsa_circ_0003428 | 4.818 | 1.466 | 0.0275085 |
| hsa_circ_0110084 | 3.570 | 1.970 | 0.0202975 |
| hsa_circ_0002346 | 6.275 | -1.356 | 0.0208140 |
| chr3\|172251260\|172262083 | 4.224 | -1.453 | 0.0189540 |
| hsa_circ_0064555 | 1.160 | 3.584 | 0.0249087 |
| hsa_circ_0066535 | 3.861 | -1.592 | 0.0361648 |
| chr4\|169507037\|169537911 | 1.108 | 3.525 | 0.0414728 |
| hsa_circ_0069562 | 1.354 | -3.663 | 0.0187866 |
| hsa_circ_0001512 | 1.479 | -2.918 | 0.0412638 |
| hsa_circ_0008321 | 1.452 | -3.059 | 0.0498562 |
| hsa_circ_0004674 | 1.082 | 3.594 | 0.0282716 |
| hsa_circ_0001722 | 8.017 | 1.137 | 0.0188014 |
| hsa_circ_0006950 | 5.259 | 1.969 | 0.0055394 |

3. List of differentially regulated circRNA identified by find_circ - tumor samples compared to normal samples

| CircRNA | baseMean | log2FoldChange | pvalue |
|---|---|---|---|
| hsa_circ_0000099 | 1.845 | 3.885 | 0.0467260 |
| hsa_circ_0000117 | 15.831 | 1.047 | 0.0103794 |
| hsa_circ_0003247 | 19.370 | 1.086 | 0.0080486 |
| hsa_circ_0000006 | 5.165 | -1.732 | 0.0497774 |
| hsa_circ_0008225 | 4.062 | 2.013 | 0.0271947 |
| hsa_circ_0006408 | 1.359 | 3.705 | 0.0377465 |
| hsa_circ_0008679 | 3.065 | 5.286 | 0.0002939 |
| hsa_circ_0006743 | 3.246 | 4.449 | 0.0017980 |
| hsa_circ_0000211 | 28.682 | 1.152 | 0.0075188 |
| hsa_circ_0003908 | 1.054 | -4.366 | 0.0257565 |
| hsa_circ_0023988 | 3.115 | 5.216 | 0.0002225 |
| hsa_circ_0023990 | 4.079 | 2.847 | 0.0154140 |
| hsa_circ_0025765 | 3.438 | -1.958 | 0.0227127 |
| chr12\|6510279\|6510532 | 1.875 | 4.125 | 0.0318281 |
| chr12\|94169153\|94186473 | 14.177 | 1.136 | 0.0479117 |
| hsa_circ_0008776 | 4.445 | -2.103 | 0.0110140 |
| hsa_circ_0002917 | 1.484 | -4.536 | 0.0176376 |
| hsa_circ_0033475 | 1.597 | -4.105 | 0.0135173 |
| hsa_circ_0000513 | 1.030 | -4.742 | 0.0160365 |
| hsa_circ_0000639 | 2.641 | 2.612 | 0.0482842 |
| hsa_circ_0039398 | 1.065 | -4.097 | 0.0199623 |
| chr17\|40495987\|40496520 | 1.544 | -4.746 | 0.0007570 |
| hsa_circ_0004621 | 1.623 | 3.812 | 0.0280436 |
| hsa_circ_0002496 | 1.824 | 4.102 | 0.0158883 |
| hsa_circ_0047136 | 1.852 | -2.168 | 0.0462213 |
| hsa_circ_0000852 | 4.280 | 2.133 | 0.0289601 |
| hsa_circ_0000915 | 18.404 | -1.406 | 0.0163990 |
| chr19\|39877663\|39893527 | 7.240 | -7.549 | 0.0188354 |
| hsa_circ_0000944 | 10.671 | -1.385 | 0.0103280 |
| hsa_circ_0110084 | 3.570 | 1.931 | 0.0441618 |
| hsa_circ_0001064 | 2.051 | -2.090 | 0.0472716 |
| hsa_circ_0001073 | 22.648 | -1.323 | 0.0146609 |
| hsa_circ_0006934 | 1.512 | 3.813 | 0.0290238 |
| hsa_circ_0002346 | 6.275 | -1.847 | 0.0033163 |
| hsa_circ_0007386 | 3.918 | -1.647 | 0.0362434 |
| hsa_circ_0002805 | 8.214 | -1.322 | 0.0273972 |
| hsa_circ_0001178 | 15.056 | 1.052 | 0.0077918 |
| hsa_circ_0001222 | 1.039 | -3.845 | 0.0303975 |
| hsa_circ_0006884 | 5.257 | -1.449 | 0.0165541 |
| hsa_circ_0006999 | 1.589 | 3.830 | 0.0274846 |
| hsa_circ_0001290 | 6.082 | -1.254 | 0.0395092 |

| | | | |
|---|---|---|---|
| hsa_circ_0001400 | 16.690 | -1.359 | 0.0016425 |
| hsa_circ_0069562 | 1.354 | -3.923 | 0.0217971 |
| chr4\|90723914\|90727297 | 1.426 | 3.593 | 0.0457490 |
| hsa_circ_0001519 | 8.444 | -1.343 | 0.0460038 |
| chr5\|123532569\|123557564 | 1.413 | 3.774 | 0.0278892 |
| hsa_circ_0001512 | 1.479 | -4.698 | 0.0012313 |
| hsa_circ_0075796 | 13.540 | 1.066 | 0.0085325 |
| hsa_circ_0001574 | 1.281 | 3.927 | 0.0419038 |
| hsa_circ_0001730 | 10.605 | -1.314 | 0.0066946 |
| hsa_circ_0004069 | 7.548 | -2.014 | 0.0033564 |
| hsa_circ_0004779 | 2.265 | 4.306 | 0.0116537 |
| hsa_circ_0079284 | 1.771 | 4.230 | 0.0093553 |
| hsa_circ_0008345 | 3.819 | -1.638 | 0.0444296 |
| hsa_circ_0008016 | 18.060 | -1.076 | 0.0317460 |
| chr8\|47394978\|47407961 | 1.233 | 4.160 | 0.0442084 |
| hsa_circ_0084582 | 1.186 | -4.211 | 0.0136427 |
| chr8\|70213903\|70216764 | 13.331 | 1.135 | 0.0154941 |
| hsa_circ_0001946 | 70.775 | 1.472 | 0.0340111 |
| hsa_circ_0009069 | 3.451 | -1.432 | 0.0470830 |

4. List of differentially regulated circRNA identified by find_circ - matched normal samples compared to normal samples

| CircRNA | baseMean | log2FoldChange | pvalue |
|---|---|---|---|
| hsa_circ_0000099 | 1.845 | 4.028 | 0.0393666 |
| hsa_circ_0000117 | 15.831 | 1.111 | 0.0070438 |
| hsa_circ_0000006 | 5.165 | -2.337 | 0.0121042 |
| hsa_circ_0002669 | 8.298 | -1.003 | 0.0350574 |
| hsa_circ_0008311 | 7.731 | 1.464 | 0.0206178 |
| hsa_circ_0008679 | 3.065 | 3.680 | 0.0151071 |
| hsa_circ_0006743 | 3.246 | 5.052 | 0.0003637 |
| hsa_circ_0019223 | 2.070 | -3.367 | 0.0063608 |
| chr11\|103219915\|103223086 | 1.575 | -2.721 | 0.0461130 |
| hsa_circ_0007401 | 1.247 | -3.565 | 0.0247624 |
| hsa_circ_0023988 | 3.115 | 3.760 | 0.0102008 |
| chr12\|28255581\|28307749 | 1.461 | -3.920 | 0.0107707 |
| chr12\|6510279\|6510532 | 1.875 | 3.912 | 0.0431521 |
| hsa_circ_0000418 | 3.027 | -2.189 | 0.0413553 |
| hsa_circ_0007881 | 2.406 | 2.715 | 0.0365187 |
| hsa_circ_0002553 | 1.727 | -2.908 | 0.0460824 |
| hsa_circ_0000513 | 1.030 | -4.455 | 0.0236707 |
| hsa_circ_0031446 | 2.450 | -2.453 | 0.0271888 |
| hsa_circ_0004621 | 1.623 | 3.832 | 0.0279417 |
| hsa_circ_0002496 | 1.824 | 3.920 | 0.0222468 |
| hsa_circ_0047720 | 1.820 | -2.917 | 0.0396129 |
| chr19\|39877663\|39893527 | 7.240 | -7.262 | 0.0238505 |
| hsa_circ_0001064 | 2.051 | -3.726 | 0.0036632 |
| chr2\|164692221\|164695836 | 1.385 | 3.984 | 0.0485440 |
| hsa_circ_0001097 | 4.017 | -1.708 | 0.0190882 |
| hsa_circ_0006934 | 1.512 | 3.671 | 0.0371467 |
| hsa_circ_0008959 | 1.100 | -4.299 | 0.0203251 |
| hsa_circ_0001178 | 15.056 | 1.008 | 0.0121514 |
| hsa_circ_0007897 | 3.058 | -2.235 | 0.0367020 |
| hsa_circ_0006884 | 5.257 | -1.555 | 0.0137264 |
| hsa_circ_0006999 | 1.589 | 3.811 | 0.0291457 |
| hsa_circ_0001290 | 6.082 | -1.312 | 0.0364634 |
| hsa_circ_0007761 | 5.638 | -1.483 | 0.0298829 |
| hsa_circ_0001400 | 16.690 | -1.349 | 0.0020922 |
| hsa_circ_0007308 | 3.100 | -2.234 | 0.0367855 |
| chr4\|90723914\|90727297 | 1.426 | 3.707 | 0.0398185 |
| hsa_circ_0001519 | 8.444 | -1.551 | 0.0243919 |
| chr5\|123532569\|123557564 | 1.413 | 3.560 | 0.0402213 |
| hsa_circ_0005524 | 1.708 | 3.997 | 0.0303311 |
| hsa_circ_0075796 | 13.540 | 1.018 | 0.0136742 |

| | | | |
|---|---|---|---|
| hsa_circ_0001571 | 2.968 | -2.205 | 0.0454621 |
| hsa_circ_0001730 | 10.605 | -1.586 | 0.0017488 |
| hsa_circ_0004779 | 2.265 | 4.298 | 0.0121305 |
| hsa_circ_0079284 | 1.771 | 3.478 | 0.0369177 |
| hsa_circ_0007672 | 2.402 | -3.348 | 0.0092848 |
| hsa_circ_0006950 | 5.259 | -1.907 | 0.0148301 |
| chr8\|70213903\|70216764 | 13.331 | 1.102 | 0.0202208 |
| hsa_circ_0008192 | 1.875 | 4.230 | 0.0286794 |
| chr9\|35657821\|35658020 | 1.525 | -4.392 | 0.0231295 |
| chrX\|1389491\|1389616 | 0.998 | -3.706 | 0.0394690 |
| hsa_circ_0001946 | 70.775 | 1.693 | 0.0147849 |
| hsa_circ_0009069 | 3.451 | -1.818 | 0.0186930 |
