## Supplementary Table 1 and 2 for "RNA sequencing identifies dysregulated circular RNAs in early-stage breast cancer"

1. List of differentially regulated circRNA identified by DCC - tumor -matched normal samples compared to normal samples

| CircRNA | baseMean | log2FoldChange | pvalue |
|---|---|---|---|
| hsa_circ_0000117 | 35.308 | 1.131 | 0.0032235 |
| chr1:149655699-149669310 | 17.897 | 1.458 | 0.0497015 |
| hsa_circ_0008339 | 8.389 | 1.316 | 0.0449583 |
| hsa_circ_0004996 | 12.846 | 1.099 | 0.0256452 |
| hsa_circ_0000175 | 10.199 | 2.417 | 0.0005345 |
| hsa_circ_0017348 | 7.318 | 1.637 | 0.0191535 |
| hsa_circ_0000032 | 7.127 | 1.394 | 0.0293706 |
| hsa_circ_0008311 | 12.077 | 1.133 | 0.0313846 |
| hsa_circ_0007503 | 3.200 | 2.023 | 0.0455362 |
| hsa_circ_0006743 | 3.106 | 2.182 | 0.0261415 |
| hsa_circ_0024605 | 7.977 | 2.507 | 0.0007389 |
| hsa_circ_0000283 | 4.488 | 1.651 | 0.0340107 |
| hsa_circ_0023984 | 9.492 | 2.890 | 0.0119991 |
| hsa_circ_0023990 | 6.716 | 1.718 | 0.0355459 |
| hsa_circ_0025765 | 7.402 | -1.594 | 0.0073208 |
| hsa_circ_0004650 | 8.180 | 1.507 | 0.0189950 |
| chr12:95208843-95211267 | 10.812 | 1.526 | 0.0399455 |
| hsa_circ_0000475 | 3.852 | 1.648 | 0.0400164 |
| hsa_circ_0000569 | 6.512 | 1.869 | 0.0047966 |
| hsa_circ_0002553 | 4.927 | -1.382 | 0.0469631 |
| hsa_circ_0009140 | 6.201 | -1.245 | 0.0368593 |
| hsa_circ_0008568 | 3.884 | 1.859 | 0.0229165 |
| chr14:49709159-49743805 | 4.769 | -1.749 | 0.0244640 |
| hsa_circ_0007695 | 23.031 | 1.513 | 0.0012597 |
| chr15:100330889-100334180 | 10.395 | 1.508 | 0.0304268 |
| hsa_circ_0007489 | 5.702 | 1.931 | 0.0161915 |
| chr15:75859878-75873568 | 8.694 | 1.380 | 0.0288581 |
| hsa_circ_0000639 | 4.855 | 2.019 | 0.0142678 |
| hsa_circ_0038005 | 5.323 | 1.897 | 0.0278916 |
| hsa_circ_0003704 | 3.518 | -1.322 | 0.0472647 |
| hsa_circ_0007068 | 7.493 | 1.645 | 0.0201046 |
| hsa_circ_0007079 | 6.174 | 2.499 | 0.0032941 |
| hsa_circ_0039943 | 13.644 | 1.747 | 0.0346147 |
| hsa_circ_0000754 | 4.079 | 1.590 | 0.0410257 |
| hsa_circ_0006220 | 5.361 | 2.316 | 0.0086691 |
| hsa_circ_0043898 | 6.016 | 1.960 | 0.0132149 |
| hsa_circ_0044623 | 3.333 | 1.697 | 0.0464428 |
| hsa_circ_0002496 | 3.582 | 2.282 | 0.0241362 |
| hsa_circ_0047880 | 4.414 | -1.524 | 0.0184299 |
| hsa_circ_0007342 | 5.365 | 2.012 | 0.0126970 |
| hsa_circ_0004321 | 12.063 | 1.563 | 0.0080331 |
| hsa_circ_0002882 | 9.873 | 1.075 | 0.0305493 |

| | | | |
|---|---|---|---|
| hsa_circ_0000963 | 22.375 | 1.871 | 0.0313819 |
| hsa_circ_0001062 | 25.818 | 2.732 | 0.0005760 |
| hsa_circ_0001064 | 8.773 | -1.174 | 0.0112565 |
| hsa_circ_0001085 | 9.008 | -1.342 | 0.0045181 |
| hsa_circ_0001097 | 6.062 | -1.367 | 0.0120988 |
| hsa_circ_0002346 | 11.267 | -1.623 | 0.0001590 |
| hsa_circ_0007386 | 9.639 | -1.143 | 0.0310433 |
| hsa_circ_0054211 | 4.496 | 2.143 | 0.0218037 |
| hsa_circ_0055451 | 5.017 | 1.590 | 0.0216843 |
| hsa_circ_0059490 | 5.764 | 2.085 | 0.0094888 |
| hsa_circ_0001187 | 18.945 | 1.359 | 0.0141584 |
| hsa_circ_0001190 | 15.059 | 1.782 | 0.0106158 |
| hsa_circ_0009055 | 4.876 | -1.351 | 0.0122604 |
| hsa_circ_0007897 | 7.824 | -1.173 | 0.0294985 |
| hsa_circ_0006884 | 9.136 | -1.167 | 0.0105129 |
| hsa_circ_0001333 | 12.169 | 1.305 | 0.0314408 |
| chr3:127014182-127014375 | 3.664 | -1.524 | 0.0379613 |
| hsa_circ_0005768 | 19.599 | 1.210 | 0.0456655 |
| hsa_circ_0008439 | 4.283 | 1.850 | 0.0220136 |
| hsa_circ_0004089 | 8.221 | -1.076 | 0.0409480 |
| hsa_circ_0007291 | 7.731 | -1.269 | 0.0173079 |
| hsa_circ_0001305 | 16.005 | 1.099 | 0.0302097 |
| hsa_circ_0066536 | 5.670 | -1.265 | 0.0374234 |
| chr4:173384651-173403950 | 4.274 | 1.546 | 0.0391134 |
| hsa_circ_0001400 | 53.016 | -1.247 | 0.0012345 |
| hsa_circ_0001410 | 7.773 | 2.193 | 0.0031509 |
| hsa_circ_0007883 | 9.578 | 2.026 | 0.0121331 |
| hsa_circ_0001519 | 16.756 | -1.238 | 0.0077342 |
| hsa_circ_0005728 | 6.985 | 1.456 | 0.0284250 |
| hsa_circ_0006716 | 5.490 | 1.462 | 0.0224033 |
| hsa_circ_0001492 | 17.766 | 1.177 | 0.0155199 |
| hsa_circ_0007132 | 10.088 | 1.572 | 0.0397623 |
| hsa_circ_0002492 | 5.955 | 1.376 | 0.0267814 |
| hsa_circ_0076125 | 5.592 | -1.334 | 0.0078957 |
| hsa_circ_0002947 | 3.005 | 1.803 | 0.0447541 |
| hsa_circ_0004069 | 12.798 | -1.304 | 0.0299554 |
| chr7:22308339-22318037 | 11.260 | 1.298 | 0.0236741 |
| chr7:30550636-30574881 | 4.470 | 1.831 | 0.0239470 |
| hsa_circ_0001829 | 48.393 | -1.041 | 0.0229182 |
| hsa_circ_0083619 | 3.412 | 2.040 | 0.0206370 |
| hsa_circ_0001790 | 7.617 | -1.316 | 0.0310913 |
| hsa_circ_0008207 | 4.958 | -1.749 | 0.0092529 |
| hsa_circ_0001946 | 221.045 | 2.098 | 0.0064935 |
| hsa_circ_0001922 | 5.700 | 1.868 | 0.0097156 |
| hsa_circ_0009069 | 6.331 | -1.065 | 0.0468476 |

2. List of differentially regulated circRNA identified by DCC - tumor samples compared to matched normal samples

| CircRNA | baseMean | log2FoldChange | pvalue |
|---|---|---|---|
| chr1:149655699-149669310 | 17.897 | 1.385 | 0.0226649 |
| hsa_circ_0003247 | 31.109 | 1.021 | 0.0150981 |
| hsa_circ_0016600 | 4.735 | 1.901 | 0.0059459 |
| hsa_circ_0016601 | 7.639 | 3.575 | 0.0000136 |
| hsa_circ_0017241 | 8.426 | 1.074 | 0.0344548 |
| hsa_circ_0009076 | 12.086 | -1.223 | 0.0336493 |
| hsa_circ_0006622 | 12.293 | -1.427 | 0.0293511 |
| hsa_circ_0008225 | 6.027 | 2.188 | 0.0018049 |
| chr10:37141718-37142290 | 19.412 | 4.060 | 0.0194411 |
| hsa_circ_0006286 | 3.941 | -1.677 | 0.0497843 |
| chr13:32535769-32537027 | 5.069 | 1.297 | 0.0383557 |
| hsa_circ_0005857 | 5.850 | 2.133 | 0.0140176 |
| hsa_circ_0002839 | 3.891 | 1.343 | 0.0451618 |
| hsa_circ_0007766 | 20.847 | 3.141 | 0.0003250 |
| hsa_circ_0003423 | 15.446 | 1.308 | 0.0087631 |
| hsa_circ_0000907 | 11.154 | 1.465 | 0.0159440 |
| hsa_circ_0059490 | 5.764 | 1.125 | 0.0410561 |
| chr20:48690947-48692790 | 5.678 | 1.606 | 0.0353722 |
| hsa_circ_0006731 | 6.432 | -1.282 | 0.0390980 |
| hsa_circ_0002174 | 18.673 | -1.582 | 0.0231743 |
| chr5:16779545-16783469 | 5.944 | -1.447 | 0.0413391 |
| hsa_circ_0005777 | 10.390 | -1.470 | 0.0029728 |
| hsa_circ_0008833 | 4.729 | 1.919 | 0.0226755 |
| chr6:19113307-19114283 | 7.882 | -1.691 | 0.0143619 |
| chr7:116110708-116112038 | 4.028 | 2.334 | 0.0035465 |
| hsa_circ_0085363 | 11.291 | 1.677 | 0.0150780 |
| hsa_circ_0006950 | 7.825 | 1.373 | 0.0440740 |
| hsa_circ_0006376 | 6.280 | -1.149 | 0.0460205 |
| hsa_circ_0008599 | 14.297 | -1.746 | 0.0062009 |
| hsa_circ_0001910 | 7.472 | 1.240 | 0.0368241 |
| hsa_circ_0001944 | 6.551 | -1.726 | 0.0065210 |
| chrX:94525211-94529483 | 7.719 | 3.246 | 0.0009262 |

3. List of differentially regulated circRNA identified by DCC - tumor samples compared to normal samples

| CircRNA | baseMean | log2FoldChange | pvalue |
|---|---|---|---|
| hsa_circ_0000117 | 35.308 | 1.115 | 0.0121146 |
| chr1:149655699-149669310 | 17.897 | 1.969 | 0.0071801 |
| hsa_circ_0003247 | 31.109 | 1.186 | 0.0166560 |
| hsa_circ_0008339 | 8.389 | 1.682 | 0.0144853 |
| hsa_circ_0004996 | 12.846 | 1.223 | 0.0312884 |
| hsa_circ_0000175 | 10.199 | 2.426 | 0.0014265 |
| hsa_circ_0016601 | 7.639 | 1.611 | 0.0384235 |
| hsa_circ_0002922 | 31.962 | 1.354 | 0.0456055 |
| chr1:247156406-247159813 | 39.615 | 1.167 | 0.0365446 |
| hsa_circ_0017348 | 7.318 | 1.961 | 0.0085074 |
| hsa_circ_0008311 | 12.077 | 1.218 | 0.0395436 |
| hsa_circ_0008225 | 6.027 | 1.958 | 0.0155203 |
| hsa_circ_0024604 | 8.465 | 1.665 | 0.0422044 |
| hsa_circ_0024605 | 7.977 | 2.245 | 0.0051530 |
| hsa_circ_0023984 | 9.492 | 3.233 | 0.0284203 |
| hsa_circ_0023990 | 6.716 | 1.994 | 0.0237407 |
| hsa_circ_0007723 | 10.776 | 1.300 | 0.0353605 |
| hsa_circ_0008250 | 5.031 | 1.613 | 0.0341010 |
| hsa_circ_0025765 | 7.402 | -2.014 | 0.0050730 |
| chr12:95208843-95211267 | 10.812 | 1.894 | 0.0169281 |
| hsa_circ_0000569 | 6.512 | 1.748 | 0.0184887 |
| chr14:33215182-33215587 | 5.385 | -1.474 | 0.0446458 |
| hsa_circ_0008568 | 3.884 | 1.802 | 0.0451523 |
| chr14:49709159-49743805 | 4.769 | -1.795 | 0.0470210 |
| hsa_circ_0007695 | 23.031 | 1.667 | 0.0013893 |
| hsa_circ_0032649 | 7.618 | -1.210 | 0.0327630 |
| chr15:100330889-100334180 | 10.395 | 1.866 | 0.0117069 |
| hsa_circ_0000596 | 13.661 | 1.053 | 0.0466980 |
| hsa_circ_0000639 | 4.855 | 2.047 | 0.0242101 |
| hsa_circ_0038005 | 5.323 | 2.092 | 0.0248642 |
| hsa_circ_0005941 | 32.773 | -1.216 | 0.0448396 |
| hsa_circ_0007068 | 7.493 | 1.601 | 0.0434424 |
| hsa_circ_0007079 | 6.174 | 2.578 | 0.0046016 |
| hsa_circ_0039943 | 13.644 | 1.961 | 0.0304504 |
| hsa_circ_0000754 | 4.079 | 1.793 | 0.0338452 |
| hsa_circ_0006220 | 5.361 | 2.118 | 0.0269304 |
| hsa_circ_0007766 | 20.847 | 2.411 | 0.0156219 |
| hsa_circ_0043898 | 6.016 | 2.017 | 0.0208728 |
| hsa_circ_0002496 | 3.582 | 2.423 | 0.0270600 |
| hsa_circ_0047880 | 4.414 | -1.529 | 0.0450529 |

| | | | |
|---|---|---|---|
| hsa_circ_0007342 | 5.365 | 2.006 | 0.0244616 |
| hsa_circ_0004321 | 12.063 | 1.362 | 0.0367715 |
| hsa_circ_0002882 | 9.873 | 1.182 | 0.0364740 |
| hsa_circ_0001062 | 25.818 | 2.961 | 0.0006005 |
| hsa_circ_0008278 | 13.240 | 1.322 | 0.0325976 |
| hsa_circ_0001073 | 49.124 | -1.556 | 0.0137440 |
| hsa_circ_0002024 | 23.569 | -1.169 | 0.0071973 |
| hsa_circ_0001085 | 9.008 | -1.309 | 0.0215865 |
| hsa_circ_0001097 | 6.062 | -1.895 | 0.0035269 |
| hsa_circ_0058040 | 8.651 | -1.487 | 0.0353461 |
| hsa_circ_0002346 | 11.267 | -2.101 | 0.0000146 |
| hsa_circ_0007386 | 9.639 | -1.715 | 0.0039812 |
| hsa_circ_0055451 | 5.017 | 1.607 | 0.0350750 |
| hsa_circ_0059490 | 5.764 | 2.548 | 0.0017228 |
| hsa_circ_0001187 | 18.945 | 1.608 | 0.0072233 |
| hsa_circ_0001190 | 15.059 | 1.751 | 0.0244057 |
| hsa_circ_0009055 | 4.876 | -1.554 | 0.0171367 |
| hsa_circ_0007897 | 7.824 | -1.324 | 0.0371722 |
| hsa_circ_0006884 | 9.136 | -1.116 | 0.0390402 |
| chr3:127014182-127014375 | 3.664 | -1.857 | 0.0326819 |
| hsa_circ_0006731 | 6.432 | -1.811 | 0.0082421 |
| chr3:151581539-151600951 | 10.411 | -1.874 | 0.0129164 |
| chr3:172251260-172262083 | 6.027 | -1.536 | 0.0123015 |
| hsa_circ_0002174 | 18.673 | -2.196 | 0.0055443 |
| hsa_circ_0008439 | 4.283 | 2.007 | 0.0232611 |
| hsa_circ_0004089 | 8.221 | -1.326 | 0.0324617 |
| hsa_circ_0071037 | 9.094 | -1.778 | 0.0265193 |
| chr4:173384651-173403950 | 4.274 | 1.659 | 0.0428116 |
| hsa_circ_0001400 | 53.016 | -1.203 | 0.0076401 |
| hsa_circ_0001410 | 7.773 | 2.142 | 0.0082077 |
| hsa_circ_0003716 | 5.294 | 1.789 | 0.0434344 |
| hsa_circ_0001519 | 16.756 | -1.250 | 0.0212755 |
| hsa_circ_0005728 | 6.985 | 1.671 | 0.0208653 |
| hsa_circ_0006716 | 5.490 | 1.457 | 0.0406390 |
| hsa_circ_0001566 | 7.803 | 1.271 | 0.0432734 |
| hsa_circ_0001492 | 17.766 | 1.390 | 0.0089206 |
| hsa_circ_0006411 | 8.433 | -1.780 | 0.0286150 |
| hsa_circ_0007132 | 10.088 | 1.778 | 0.0348825 |
| hsa_circ_0076125 | 5.592 | -1.600 | 0.0081159 |
| hsa_circ_0004069 | 12.798 | -1.718 | 0.0104304 |
| chr7:30550636-30574881 | 4.470 | 2.079 | 0.0168490 |
| hsa_circ_0002714 | 7.431 | -1.588 | 0.0449351 |
| hsa_circ_0001721 | 5.167 | 1.629 | 0.0337511 |
| hsa_circ_0008345 | 6.341 | -1.289 | 0.0318086 |
| hsa_circ_0001829 | 48.393 | -1.459 | 0.0026867 |

| | | | |
|---|---|---|---|
| hsa_circ_0083619 | 3.412 | 2.066 | 0.0316977 |
| hsa_circ_0008207 | 4.958 | -1.663 | 0.0344018 |
| hsa_circ_0001944 | 6.551 | -1.848 | 0.0094817 |
| hsa_circ_0001946 | 221.045 | 2.001 | 0.0200872 |
| hsa_circ_0001922 | 5.700 | 1.543 | 0.0491799 |

4. List of differentially regulated circRNA identified by DCC - matched normal samples compared to normal samples

| CircRNA | baseMean | log2FoldChange | pvalue |
|---|---|---|---|
| hsa_circ_0000117 | 35.308 | 1.115 | 0.0121146 |
| chr1:149655699-149669310 | 17.897 | 1.969 | 0.0071800 |
| hsa_circ_0003247 | 31.109 | 1.186 | 0.0166559 |
| hsa_circ_0008339 | 8.389 | 1.682 | 0.0144852 |
| hsa_circ_0004996 | 12.846 | 1.223 | 0.0312883 |
| hsa_circ_0000175 | 10.199 | 2.426 | 0.0014265 |
| hsa_circ_0016601 | 7.639 | 1.611 | 0.0384231 |
| hsa_circ_0002922 | 31.962 | 1.354 | 0.0456051 |
| chr1:247156406-247159813 | 39.615 | 1.167 | 0.0365443 |
| hsa_circ_0017348 | 7.318 | 1.961 | 0.0085073 |
| hsa_circ_0008311 | 12.077 | 1.218 | 0.0395434 |
| hsa_circ_0008225 | 6.027 | 1.958 | 0.0155201 |
| hsa_circ_0024604 | 8.465 | 1.665 | 0.0422041 |
| hsa_circ_0024605 | 7.977 | 2.245 | 0.0051530 |
| hsa_circ_0023984 | 9.492 | 3.233 | 0.0284200 |
| hsa_circ_0023990 | 6.716 | 1.994 | 0.0237405 |
| hsa_circ_0007723 | 10.776 | 1.300 | 0.0353603 |
| hsa_circ_0008250 | 5.031 | 1.613 | 0.0341008 |
| hsa_circ_0025765 | 7.402 | -2.014 | 0.0050730 |
| chr12:95208843-95211267 | 10.812 | 1.894 | 0.0169280 |
| hsa_circ_0000569 | 6.512 | 1.748 | 0.0184886 |
| chr14:33215182-33215587 | 5.385 | -1.474 | 0.0446459 |
| hsa_circ_0008568 | 3.884 | 1.802 | 0.0451522 |
| chr14:49709159-49743805 | 4.769 | -1.795 | 0.0470213 |
| hsa_circ_0007695 | 23.031 | 1.667 | 0.0013893 |
| hsa_circ_0032649 | 7.618 | -1.210 | 0.0327631 |
| chr15:100330889-100334180 | 10.395 | 1.866 | 0.0117069 |
| hsa_circ_0000596 | 13.661 | 1.053 | 0.0466977 |
| hsa_circ_0000639 | 4.855 | 2.047 | 0.0242100 |
| hsa_circ_0038005 | 5.323 | 2.092 | 0.0248641 |
| hsa_circ_0005941 | 32.773 | -1.216 | 0.0448399 |
| hsa_circ_0007068 | 7.493 | 1.601 | 0.0434422 |
| hsa_circ_0007079 | 6.174 | 2.578 | 0.0046015 |
| hsa_circ_0039943 | 13.644 | 1.961 | 0.0304502 |
| hsa_circ_0000754 | 4.079 | 1.793 | 0.0338450 |
| hsa_circ_0006220 | 5.361 | 2.118 | 0.0269302 |
| hsa_circ_0007766 | 20.847 | 2.411 | 0.0156216 |
| hsa_circ_0043898 | 6.016 | 2.017 | 0.0208727 |
| hsa_circ_0002496 | 3.582 | 2.423 | 0.0270599 |
| hsa_circ_0047880 | 4.414 | -1.529 | 0.0450532 |
| hsa_circ_0007342 | 5.365 | 2.006 | 0.0244615 |

| | | | |
|---|---|---|---|
| hsa_circ_0004321 | 12.063 | 1.362 | 0.0367713 |
| hsa_circ_0002882 | 9.873 | 1.182 | 0.0364738 |
| hsa_circ_0001062 | 25.818 | 2.961 | 0.0006005 |
| hsa_circ_0008278 | 13.240 | 1.322 | 0.0325974 |
| hsa_circ_0001073 | 49.124 | -1.556 | 0.0137441 |
| hsa_circ_0002024 | 23.569 | -1.169 | 0.0071974 |
| hsa_circ_0001085 | 9.008 | -1.309 | 0.0215867 |
| hsa_circ_0001097 | 6.062 | -1.895 | 0.0035269 |
| hsa_circ_0058040 | 8.651 | -1.487 | 0.0353463 |
| hsa_circ_0002346 | 11.267 | -2.101 | 0.0000146 |
| hsa_circ_0007386 | 9.639 | -1.715 | 0.0039812 |
| hsa_circ_0055451 | 5.017 | 1.607 | 0.0350749 |
| hsa_circ_0059490 | 5.764 | 2.548 | 0.0017228 |
| hsa_circ_0001187 | 18.945 | 1.608 | 0.0072232 |
| hsa_circ_0001190 | 15.059 | 1.751 | 0.0244055 |
| hsa_circ_0009055 | 4.876 | -1.554 | 0.0171367 |
| hsa_circ_0007897 | 7.824 | -1.324 | 0.0371724 |
| hsa_circ_0006884 | 9.136 | -1.116 | 0.0390404 |
| chr3:127014182-127014375 | 3.664 | -1.857 | 0.0326820 |
| hsa_circ_0006731 | 6.432 | -1.811 | 0.0082422 |
| chr3:151581539-151600951 | 10.411 | -1.874 | 0.0129165 |
| chr3:172251260-172262083 | 6.027 | -1.536 | 0.0123015 |
| hsa_circ_0002174 | 18.673 | -2.195 | 0.0055443 |
| hsa_circ_0008439 | 4.283 | 2.007 | 0.0232610 |
| hsa_circ_0004089 | 8.221 | -1.326 | 0.0324619 |
| hsa_circ_0071037 | 9.094 | -1.778 | 0.0265195 |
| chr4:173384651-173403950 | 4.274 | 1.659 | 0.0428115 |
| hsa_circ_0001400 | 53.016 | -1.203 | 0.0076402 |
| hsa_circ_0001410 | 7.773 | 2.142 | 0.0082076 |
| hsa_circ_0003716 | 5.294 | 1.789 | 0.0434341 |
| hsa_circ_0001519 | 16.756 | -1.250 | 0.0212757 |
| hsa_circ_0005728 | 6.985 | 1.671 | 0.0208652 |
| hsa_circ_0006716 | 5.490 | 1.457 | 0.0406388 |
| hsa_circ_0001566 | 7.803 | 1.271 | 0.0432732 |
| hsa_circ_0001492 | 17.766 | 1.390 | 0.0089205 |
| hsa_circ_0006411 | 8.433 | -1.780 | 0.0286151 |
| hsa_circ_0007132 | 10.088 | 1.778 | 0.0348822 |
| hsa_circ_0076125 | 5.592 | -1.600 | 0.0081160 |
| hsa_circ_0004069 | 12.798 | -1.718 | 0.0104304 |
| chr7:30550636-30574881 | 4.470 | 2.079 | 0.0168488 |
| hsa_circ_0002714 | 7.431 | -1.588 | 0.0449353 |
| hsa_circ_0001721 | 5.167 | 1.629 | 0.0337509 |
| hsa_circ_0008345 | 6.341 | -1.289 | 0.0318087 |
| hsa_circ_0001829 | 48.393 | -1.459 | 0.0026868 |
| hsa_circ_0083619 | 3.412 | 2.066 | 0.0316976 |

| | | | |
|---|---|---|---|
| hsa_circ_0008207 | 4.958 | -1.663 | 0.0344020 |
| hsa_circ_0001944 | 6.551 | -1.848 | 0.0094817 |
| hsa_circ_0001946 | 221.045 | 2.001 | 0.0200869 |
| hsa_circ_0001922 | 5.700 | 1.543 | 0.0491798 |
