## Supplementary Table 3 for "RNA sequencing identifies dysregulated circular RNAs in early-stage breast cancer"

Histopathological status of samples

| S. No. | Tissue type | Sample ID | Age | IDC GRADE | TNM staging | ER | PR | HER2 |
|---|---|---|---|---|---|---|---|---|
| 1 | Breast Tumor | T 1 | 61 | IDC G II | T1N1M0 | + | + | - |
| 2 | Breast Tumor | T 2 | 53 | IDC G II-III | T1N1M0 | + | + | + |
| 3 | Breast Tumor | T 3 | 61 | IDC G III | T1 N1 M0 | - | - | + |
| 4 | Breast Tumor | T 4 | 38 | IDC G II | T2 N0 M0 | + | + | + |
| 5 | Breast Tumor | T 5 | 46 | IDC G II | T1N0M0 | + | + | - |
| 6 | Breast Normal | N 1 | 47 | NA | NA | NA | NA | NA |
| 7 | Breast Normal | N 2 | 46 | NA | NA | NA | NA | NA |
| 8 | Breast Normal | N 3 | 50 | NA | NA | NA | NA | NA |
